# FMRI processing with AFNI: Some comments and corrections on “Exploring the Impact of Analysis Software on Task fMRI Results”

**DOI:** 10.1101/308643

**Authors:** Paul A. Taylor, Gang Chen, Daniel R. Glen, Justin K. Rajendra, Richard C. Reynolds, Robert W. Cox

## Abstract

A recent study posted on bioRxiv by Bowring, Maumet and Nichols aimed to compare results of FMRI data that had been processed with three commonly used software packages (AFNI, FSL and SPM). Their stated purpose was to use “default” settings of each software’s pipeline for task-based FMRI, and then to quantify overlaps in final clustering results and to measure similarity/dissimilarity in the final outcomes of packages. While in theory the setup sounds simple (implement each package’s defaults and compare results), practical realities make this difficult. For example, different softwares would recommend different spatial resolutions of the final data, but for the sake of comparisons, the same value must be used across all. Moreover, we would say that AFNI does not have an explicit default pipeline available: a wide diversity of datasets and study designs are acquired across the neuroimaging community, often requiring bespoke tailoring of basic processing rather than a “one-size-fits-all” pipeline. However, we do have strong recommendations for certain steps, and we are also aware that the choice of a given step might place requirements on other processing steps. Given the very clear reporting of the AFNI pipeline used in Bowring et al. paper, we take this opportunity to comment on some of these aspects of processing with AFNI here, clarifying a few mistakes therein and also offering recommendations. We provide point-by-point considerations of using AFNI’s processing pipeline design tool at the individual level, afni_proc.py, along with supplementary programs; while specifically discussed in the context of the present usage, many of these choices may serve as useful starting points for broader processing. It is our intention/hope that the user should examine data quality at every step, and we demonstrate how this is facilitated in AFNI, as well.

## INTRODUCTION

There are many choices to be made when designing and carrying out a neuroimaging study, each of which can have long-ranging consequences on the final outcome. Study design is a notable one, and there is a wide array of implementations that are considered reasonable within the FMRI community: resting, block design task, event-related task, naturalistic movie-task, feedback-tasks, etc., each of which may have differing chosen parameters (trial duration, timing gap between stimuli, randomization across trials, eyes-open or -closed rest, type of movie, with or without headphones, etc.). Additionally, researchers can choose to gather supplementary data for inclusion in the analysis, such as monitoring subjects’ heart rate, pulse, breathing, head motion, alertness and responses. Scanners themselves may vary in field strength and number of head coils, greatly affecting data contrast levels and potential distortions. In many cases the choice of experimental design data and specific acquisition parameters will closely inform the processing pipeline choices on the way to statistical analysis.

Given this plethora of data types and characteristics, the AFNI (Cox, 1996) toolbox has historically *not* had a default pipeline (in the sense of one set of steps that we *always* advocate) -- we have left the processing specification with those very same scientists who are designing and carrying out the study. That being the case, we certainly have made recommendations on individual steps and combinations of tools, and we provide these freely in program help files, in consultations, in class handouts and lectures, on the AFNI Message Board (which is publicly available) and in papers that we have published (a recent example being Cox et al. (2017a), with scripts available online^1^). As we continually update existing tools and develop new ones, even these recommendations can, and certainly do, change over time.

In a recently uploaded draft Bowring, Maumet and Nichols (2018; referred to as “BMN” hereafter^2^) used publicly available task-based FMRI data to compare results of processing pipelines. The stated goal was to look at the variability in statistical values and outcomes due to the usage of different software packages (measured in terms of differences and pairwise comparisons). For this, BMN investigated three commonly used FMRI toolboxes, AFNI, FSL (Smith et al., 2004), and SPM (Penny et al., 2011), using what they term as “default” settings in each. This task is inherently challenging. Firstly, as noted above, AFNI itself does not have a recommended default, in the sense that there is a pre-built pipeline to run without regard for the input data’s specifics (e.g., field strength, voxel size, TR, tissue contrast, subject motion characteristics). Furthermore, packages have different output preferences (for example, about upsampling spatial resolution), as well as different definitions of basic features like voxel “neighbors” (in AFNI, default is facewise, “nearest neighbor” NN=1; in SPM, face+edgewise, NN=2; in FSL, face+edge+nodewise, NN=3) and default coordinate systems (in AFNI, the default is RAI/DICOM; in SPM and FSL, it is LPI). Then there are analysis choices made by the researching scientists from within a range of generally accepted values (e.g. blur size, template, alignment method, censor limits, group masking parameters).

Therefore, we aim to review and discuss some of the processing steps in AFNI that were implemented in BMN. It must be noted that this is made possible by the fact that those authors ensured easy access to the data collections that were investigated and to the processing scripts themselves (for all softwares), and this is certainly a very laudable feature of their work. We also repeat the processing of two of the test studies carried out by BMN using the same publicly available data, and describe the processing steps we would have initially chosen if presented with this study design, along with brief comments on interpreting the output of AFNI’s statistical analyses. We compare aspects of analysis using their specified AFNI commands and the presently recommended ones. Our focus is on performing reasonable FMRI processing and how this can be accomplished with AFNI, and we do not comment on choices made for SPM or FSL, nor on those softwares’ defaults, nor on the design or final results/cluster locations of this particular FMRI study.

## METHODS AND RESULTS

The first two group analyses in the BMN paper are repeated here, using the same task-based data available from OpenfMRI (Poldrack et al., 2013). The ds000001 dataset (release 2.0.4) contains 16 healthy adults, and the ds000109 dataset (release 2.0.2) has 21 healthy adults available. In each case the group analysis is a one-sample test with mixed effects modeling^3^ carried out voxelwise, with additional cluster-based corrections for familywise errors (FWEs). The AFNI version used here is 18.0.27.

With AFNI, the data processing steps are primarily specified through the toolbox’s pipeline formulator of choice, afni_proc.py. Using the afni_proc.py command, a full processing stream is specified and tailored with options (typically, 20-40 lines of code), and a fully commented shell script is created (typically 400-500+ lines of code). Typical inputs include: datasets (anatomical, functional EPI, etc.), major processing steps (alignment, volume registration, blurring, subject level regression modeling, etc.) and options for any processing step (blur size, registration cost function, tissue regressor type, etc.). Most intermediate datasets during processing are retained, for both quantitative and visual quality control (QC) checks by the analyst. Importantly, afni_proc.py does not just create a processing script, but it also automatically generates supplementary scripts for “single subject reviews,” to facilitate checking the results of analysis carefully and efficiently.^4^ In order to evaluate group properties and outliers, the AFNI program gen_ss_review_table.py can create a single spreadsheet or table of the pertinent QC numbers from multiple subjects’ analyses that were automatically calculated and recorded during processing (quantities of motion, censoring, degrees of freedom remaining in the data, etc.). These and other supplementary programs are discussed further below.

We organize the processing comments as follows. First, we discuss aspects of BMN’s AFNI processing that *need* to be modified, for mathematical or other correctness (A). We then present features that we feel *should* be adjusted, for improved methodology (B). Finally, we comment on additional considerations or aspects of FMRI processing (C). Throughout this discussion of point-by-point analysis choices, we present some results of running: the same AFNI processing specified by BMN (hereafter, “BMN-AFNI” or simply “BMN”; though, again, we use the more current version of AFNI noted above); the BMN-AFNI processing with one additional option that must be used in 3dMEMA to avoid edge-induced artifacts; and our presently preferred processing for these datasets^5^ (“NIMH-AFNI”). In Table 1, we have replicated the breakdown of processing steps from BMN-Table 1 for the AFNI software (with some additions and some minor terminological alterations), listing the BMN-AFNI options as well as any NIMH-AFNI options that differed; a column of reference numbers is provided to refer to discussion points within the text. Finally, we also display the cluster maps of positive and negative effects from BMN-AFNI processing with those from NIMH-AFNI for ds000001 (D); we do not compare final outputs deeply, as the focus of the present work is to discuss processing steps, and there is also no “gold standard” result for comparison.

**Table 1:** The following is a list of processing steps of AFNI (following the format of Table 1 in BMN) for the implementations used by BMN (“BMN-AFNI” column) and those we have used in the present work (“NIMH-AFNI”). The final column contains a reference number, for location discussion of points within the text.

| Stage | Step | BMN-AFNI | NIMH-AFNI | Ref. |
| --- | --- | --- | --- | --- |
| Pre-preproc | Command |  | @SSwarper<br>(using a high-definition template) | B-1 |
| Preproc. | Script | via afni_proc.py |  |  |
|  | Delete first time points | -tcat_remove_first_trs 2 | [-tcat_remove_first_trs 2] | C-4 |
|  | Slice-timing | -tshift_opts_ts -tpattern |  |  |
|  | Realignment/motion correction<sup>†</sup> | -volreg_align_e2a | -volreg_align_e2a<br>-volreg_align_to MIN_OUTLIER | B-3 |
|  | Segmentation | *Not applied* |  |  |
|  | Skull stripping anat. (brain extr.) | *Implicitly run within align block* | -copy_anat [+ @SSwarper results]<br>-anat_has_skull no | B-1 |
|  | Mask func. | *Not applied* |  |  |
|  | Intrasubject coregistration (func. to anat.)<sup>†</sup> | -align_opts_aea<br>-giant_move<br>-check_flip | -align_opts_aea -ginormous_move<br>-check_flip -cost lpc+ZZ | B-2 |
|  | intersubject registration (anat. to standard) | -tlrc_base<br>-volreg_tlrc_warp | -volreg_tlrc_warp -tlrc_NL_warp<br>-tlrc_NL_warped_dsets [+ @SSwarper results] | B-1 |
|  | Analysis (final) voxel size | -volreg_warp_dxyz 2<br>(overriding default determined from functional images) | (NB: here, we upsample to match other software choices; under normal circumstances, we would **not** upsample heavily) | C-1 |
|  | Smooth/blur | -blur_size |  |  |
|  | Scaling<sup>*</sup> | *BMN included the "scale" block, but did not note this explicitly in the text.* | [The "scale" block was included.] | C-3 |
| Subject level<sup>**</sup> | Script | via afni_proc.py |  |  |
|  | Model specification | -regress_stim_times<br>-regress_stim_labels<br>-regress_basis_multi<br>-regress_stim_types |  | A-5,<br>A-6 |
|  | **Inclusion of 6 motion parameters** | *Implicitly added within 'regress' block* | *Implicitly added within 'regress' block, and also: -regress_motion_per_run* | C-6 |
|  | **Motion-based censoring\*\*** | -regress_censor_motion 0.3 | [-regress_censor_motion 0.3] | C-5 |
|  | **Model estimation** | *Nothing to specify* |  |  |
|  | **Contrasts** | -regress_opts_3dD<br>-gltsym | -regress_opts_3dD -gltsym<br>-regress_3dD_stop -regress_reml_exec | A-3 |
|  | **Single subject QC†** |  | @snapshot_volreg, @ss_review_basic, @ss_review_driver | C-7 |
| **Group level\*\*\*** | **Script** | 3dMEMA | gen_group_command.py<br>(easier to make 3dMEMA command) | C-8 |
|  | **Model specification** | 3dMEMA -set | also include in above gen*.py:<br>-options -missing_data 0 | A-1 |
|  | **Group QC†** |  | gen_ss_review_table.py | C-7 |
|  | **Contrasts** | *Nothing to specify* |  |  |
|  | **Masking\*\*\*\*** | 3dMean (obtain group-mask) | *Simplified steps-- AFNI programs can output NIFTI directly* | A-2 |
|  | **Clusterizing\*\*\*\*** | 3dClustSim |  | C-2 |
|  | **Making cluster masks and final outputs\*\*\*\*** | 3dclust, 3dcalc (Binarizing cluster masks and masking t-stat)<br>3dAFNItoNIFTI (Converting from .BRIK/.HEAD to .nii) |  | A-4,<br>C-9 |

† We note that these two processing step labels, "Realignment/motion correction" and "Intrasubject coregistration (func. to anat.)," were swapped in the BMN description; we would consider the present arrangement here to be a more accurate assignment of duty of the afni\_proc.py options listed.
‡ These processing steps were not included in the BMN list, but we feel that they should be considered part of standard processing at both the subject and group levels.
\* Even though not explicitly noted in BMN, the "scale" block was included in afni\_proc.py, which we strongly recommend and included in the NIMH-AFNI version, as well. See the text for additional comments on scaling time series.
\*\* BMN did not list this in their table and moreover stated that they did not censor volumes based on motion (see C-5 section here). However, they did, in fact, include this option and implement motion-based censoring via afni\_proc.py. We would certainly generally recommend doing so.
\*\*\* Instead of the BMN terminology "level 1" and "level 2," we use "subject level" and "group level," respectively. We note that different softwares use level numbers to refer to different stages, so we adopt descriptive labels for clarity.
\*\*\*\* We divided the many steps listed by BMN in a single section of the table "Second-level inference" into separate categories, to more easily discuss choices of the individual pieces.

### A) Steps in BMN that need to be modified

#### [A-1] Marking missing data to avoid erroneous modeling

At the group analysis stage, result maps from individual subjects’ datasets are combined. After alignment and warping of the functional EPI data to standard space, there will be zero values outside the warped volume’s field of view (FOV), and the warped FOV edges may not exactly overlap across the group. Simply leaving those (one or more) zero values of missing data in the model would lead to erroneous and meaningless results at those voxels (due to divisions by zero); such cases could be recognized as producing *t*-statistics with extreme values at or near ±100 (the default limits within AFNI statistics programs). To avoid such numerical pathology, the option “-missing_data 0” should be used when running 3dMEMA; for 3dttest++, the analogous option is “-zskip” Similarly heterogeneous masking would arise if individual subject maps had been directly applied to each subject’s results and then run in 3dMEMA or 3dttest++, and the same options should be applied.

The absence of such zero-ignoring options in BMN’s analysis is responsible for their following comments:

> *“For ds000109, the group-level activation map for parametric inference in AFNI contains a small cluster of voxels in the cerebellum near the edge of the analysis mask showing unusually large T-statistic values, with many voxels having a T-statistic of exactly 100 (a bound set by AFNI’s 3dMEMA).”*

The actual cause and solution of this “problem” is shown in Fig. 1, where we re-performed their analysis on the ds000109 dataset without (top row) and with (middle row) the correct zero-masking options. The top row shows the unmasked *t*-statistic results from 3dMEMA (à la BMN); the edges of the EPI volumes FOV do not exactly coincide in standard space, meaning that some, but not all, subjects’ values were zero there, leading to *t*-statistics of extremely large magnitude (~100, as highlighted with magenta arrows) in this case. While the same phenomenon occurs in other datasets (such as ds000001), it is most noticeable here because these warped FOV edges intersect with the brainstem of the template data, due to the method of data collection. Without applying the “-missing_data 0” option, artifactual clusters could remain in the output, as in the BMN analysis; applying the proper program option removes these results. Furthermore, as discussed in the next point, a different group mask should also be used, restricting whole brain analysis to the intersection of all subjects’ EPI and anatomical data (shown in the bottom row).

**Figure 1.**
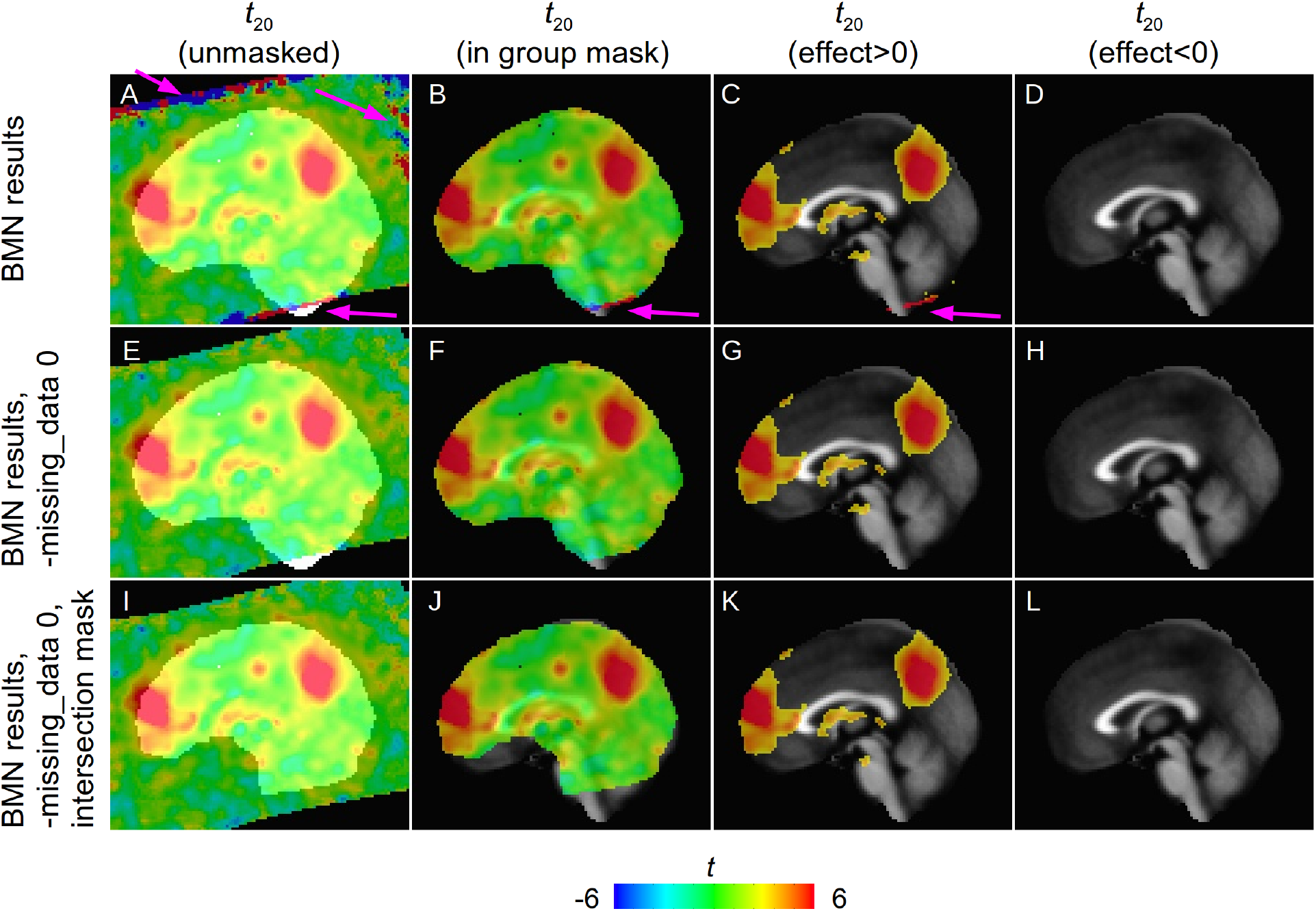
A comparison of masking and group level approaches with ds000109 (cf BMN-Fig. 1, middle column). In the top row panel A shows BMN’s unmasked *t*-statistic maps (underlay is their group mask), with some very large magnitudes present (~100, highlighted with magenta arrows), due to their inclusion of FOV edge artifacts by not masking out zero values; B shows that these edge-artifacts remain in their group mask and then C-D show these in their clusterized results (in rerunning their data, a small negative-result cluster did appear, but in a different slice). In the middle row (EH), after the same BMN-AFNI preprocessing 3dMEMA was run using the correct “-missing_data 0” option, and the artifacts are not present. In the bottom row (I-L), we display their results with an improved group-level mask, which is created from each subject’s anatomical and EPI data rather than from the template extents (discussed more in point A-2). The cluster extent thresholds (voxelwise threshold *p*=0.005, FWE=5%, one-sided test, NN=1) from top to bottom were: 482, 482 and 442. In all cases: sagittal slices; *x*=0R; image left=subject anterior.

BMN also noted that the very large statistics values contribute to the streaks observed in their Fig. 7. Again, and for the same reasons as noted above, these would be avoided with the appropriate usage of “3dMEMA -missing_data 0 …”.

#### [A-2] Making a group mask

Masking at the group level typically tries to balance between: restricting analyses to the brain volume, leaving out noise-dominated regions, and reducing the number of “multiple comparison” corrections that are required to control false positive rates (FPRs), while also not leaving out parts of the brain that may have expected activity and results.^6^ We generally recommend to leave masking until the group analysis stage, and when using afni_proc.py, one may create a group mask from the intersection of each subject’s EPI masks, with the *mask_epi_anat* dataset currently preferred.^7^ These masks are determined from the estimated brain coverage of the EPI and anatomical data for each subject, and their intersection provides a more representative mask of the extent of usable data across the group. From BMN’s level2.sh script, it appears that they used the intersection of the *mask_group* datasets, instead; this is created from the skull-stripped template image (which is the same across the group), and this can lead to the inclusion of many voxels outside the actual EPI data’s coverage (esp. in the presence of signal dropout, as in the present data; see Figs. 6-7), so that clustering results are likely overly conservative (due to the larger volume) and have a higher likelihood of containing extra-cranial “noise” clusters (due to the nature of the extra regions).

A comparison of the BMN-generated group mask and the NIMH-AFNI group mask is shown in Fig. 2, where hot colors show the relative overlap of all subject masks, and blue shows the final mask. The BMN mask contains 245,683 voxels, and the NIMH-AFNI mask contains 193,142 voxels, 79% of the former. We note that the mask volume or voxel count should likely be reported when reporting cluster values, as they affect the FWE corrections.

**Figure 2.**
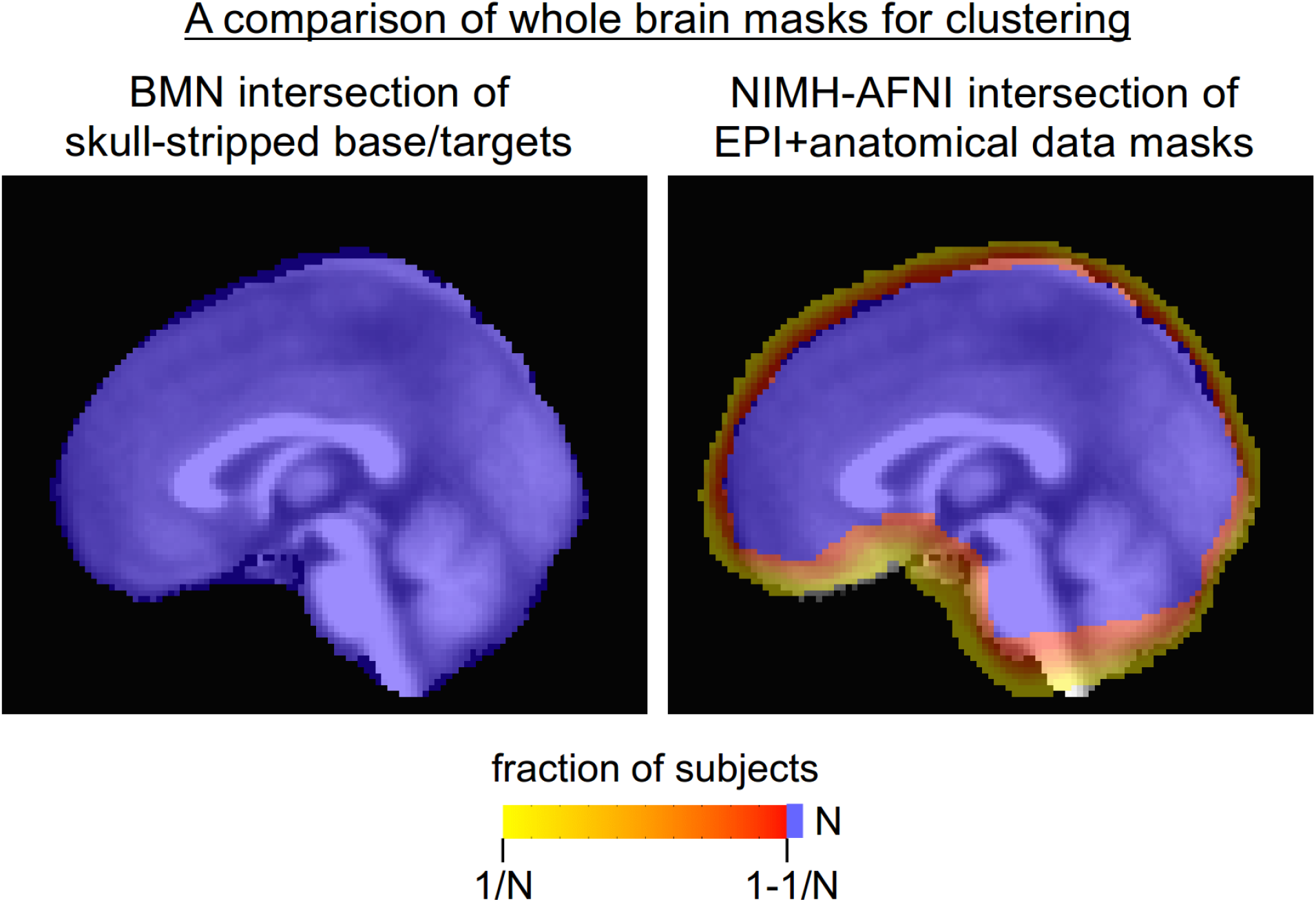
A comparison of the group level masks used for clustering, in this case with the N=21 subjects of ds000109 (sagittal slices, *x*=0, image left=subject anterior). Left: BMN took the intersection of masks determined by the template, which are constant across the group; importantly, this does not reflect the coverage of each subject’s EPI and anatomical information and would potentially include many areas of low signal. Right: the NIMH-AFNI group mask was generated by using the individual extents of EPI and anatomical data (non-total overlap in hot colors), whose intersection (blue) provides a mask of usable data across the group. Note that using the BMN mask would lead to more conservative FWE corrections, as well, due to its larger size.

Additionally, the use of the template-based mask in the BMN group analysis means that, for some subjects in the present dataset, voxels with missing data were included in the statistical modeling; and as noted above, unless “-missing_data 0” is used in 3dMEMA, or “-zskip” in 3dttest++, then results at the edge and exterior of the brain would simply be wrong.

#### [A-3] Proper modeling with 3dMEMA

In order to use 3dMEMA (i.e., to perform Mixed Effects Multilevel Analysis at the group level^8^), one *must* use 3dREMLfit instead of 3dDeconvolve in afni_proc.py. Using 3dMEMA, both the effect estimates *and* the *t*-statistics from the subject level are carried to the group level. For estimating these quantities, 3dREMLfit performs generalized least squares (GLSQ) modeling with a restricted maximum likelihood (REML) approach in order to account for temporal correlation (Chen et al., 2012), and is a generalized and more accurate approach for estimating the *t*-statistic than that of 3dDeconvolve, which uses ordinary least squares (OLSQ) modeling. The underlying assumption of 3dDeconvolve’s OLSQ approach to computing *t*-statistics -- that the residual time series is independent across time -- is often not reasonable in FMRI data, and 3dREMLfit instead models the temporal correlation structure more appropriately (Olzsowy et al., 2017). In afni_proc.py the adoption of 3dREMLfit would be done with options *“-regress_3dD_stop -regress_reml_exec.”*

We note that the effect estimates from 3dDeconvolve’s OLSQ approach are generally adequate for group level analyses in other approaches (say, for *t*-tests with 3dttest++). However, there is little reason not to use 3dREMLfit for this purpose as well. In the near future, 3dREMLfit will likely become the pre-selected time series modeler in afni_proc.py.

#### [A-4] Two-sided test vs paired one-sided tests

Here we discuss the manner in which testing is done, referring to the combination of modeling (calculating statistic values, in the present case meaning *t*-statistics from 3dMEMA) and thresholding (applying the user’s desired significance level and performing voxelwise thresholding). BMN set up their voxelwise analysis as a pair of one-sided tests, looking for positive and negative results separately. When several statistical tests are performed simultaneously, multiple comparisons correction must be utilized in order to avoid inflating the overall false positive rate. Bonferroni correction is a standard approach for this, whereby the threshold *p*-value is corrected by dividing by the number of multiple tests *M*. In the case of a pair of one-sided tests, *M*=2. Importantly, performing a two-sided test with a given *p*-value *P* is mathematically equivalent to performing a pair of one-sided tests *using* the Bonferroni corrected *p*-value of *P/2*. Not including this correction when performing such a pair of one-sided tests is incorrect and can lead to an inflated false positive rate.

If one has specific, predetermined hypothesis that an effect difference would have a certain directionality (say, *A*>0), then performing one-sided testing can be justified. Such a scenario may be particularly relevant when investigating a localized region of the brain, for example. However, if such a strong hypothesis does *not* exist a priori for the area of focus (esp. when performing whole brain comparisons), then two-sided testing must be used for correctness. One might suggest that using a pair of one-sided tests is useful for classifying clusters of “positive” effects and “negative” effects separately; however, such is not necessary, and can likely lead to incorrect thresholding. With two-sided testing one converts *p*-to-*t* appropriately to obtain a threshold *t*\*, and then the directionality of a cluster is posteriorly revealed through the sign of the effect estimate or the corresponding *t*-value.

Consider calculating the *t*-statistic threshold associated with a given *p* shown in Fig. 1. The one-sided *p*=0.01 translates to *t*_15_=2.602 (positive direction) or −2.602 (negative direction), and by simultaneously testing both directions would obtain clusters of statistic values *t*_15_>2.602 and *t*_15_<−2.602, which are shown in the third and fourth columns of that figure. In comparison, to perform equivalent clustering in a two-sided case, one finds that *p*=0.02 translates to *t*_15_=2.602, leading to the same clusters and in agreement with the Bonferroni correction equivalency noted above; using a two-sided *p*=0.01 would lead to a higher statistic threshold *t*_15_=2.946, which would not provide equivalent results.

#### [A-5] High correlations among regressor pairs

The linear regression in ds000001 was set up by BMN to use 3 regressors per stimulus condition: an average response, a parametrically modulated response, and one that varied with the pump duration. Since the onsets of the average and duration-modulated events are identical, these regressor pairs end up being highly correlated (|*r*|>0.9), making the resulting beta weights unstable and less suitable for such a comparative analysis. The user is warned of these high correlations during the afni_proc.py processing and again when running the highly recommended quality control review script, @ss_review_driver, along with plots of the regressors of interest. The warning message output by the review script for sub-01 for the BMN-AFNI processing was:

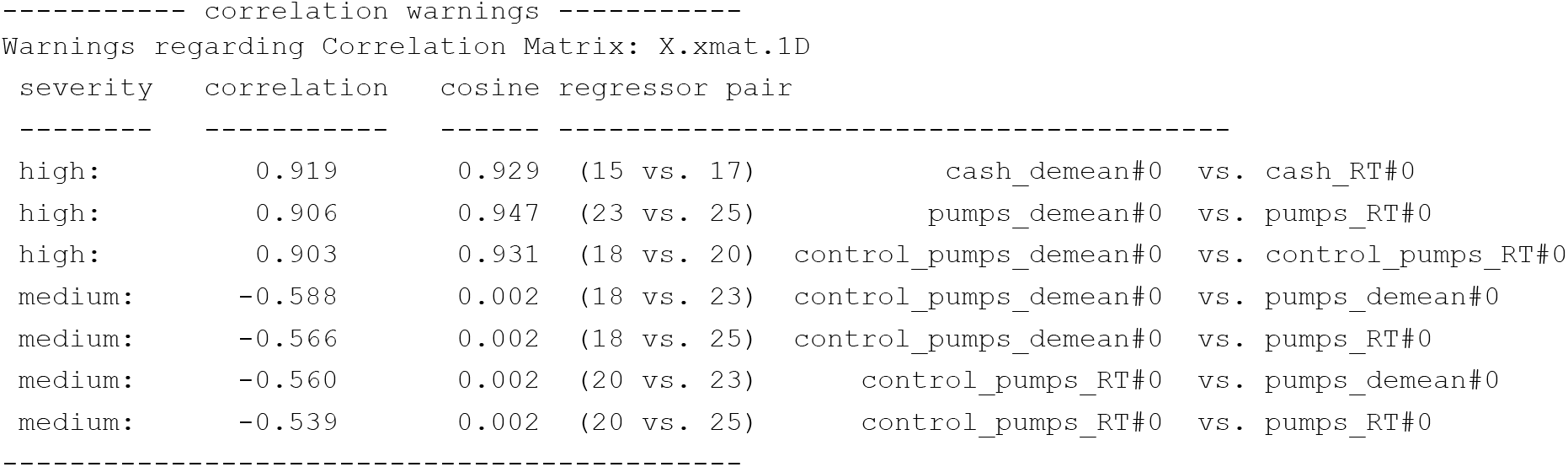

In the original analysis (Schonberg et al., 2012), the duration modulated regressors were made to be orthogonal to the average response regressors. This pre-modeling step alleviates the concern over such instability, while effectively merging the duration modulated responses into the average one. That could be done in AFNI by generating and orthogonalizing the duration-modulated regressors before giving them to afni_proc.py. Note that this pre-orthogonalization would affect the final model results (as the regressors themselves are changed) and have to be described in detail (e.g., the order in which regressors are orthogonalized changes their values).

#### [A-6] Choosing the contrast for group analysis

Based on the original analysis (Schonberg et al., 2012), it might have been more expected to run the group analysis based on the average response to pumps minus the average response to control. However, the BMN analysis used the the sum of the average and modulated betas for pumps minus the corresponding sum for control. This might have been an oversight, where ‘pumps_demean[0]-control_pumps_demean[0]’ should have been used for the general linear test in the regression.^9^ The BMN analysis includes the average response which is highly correlated with the RT regressor, as noted above, along with the modulated terms, which should be mostly uncorrelated.

### B) Points that should be adjusted

#### [B-1] Alignment to standard space

We *strongly* recommend using nonlinear registration (“warping”) for the alignment of subject anatomical volumes to standard space (“intersubject registration”). In fact, as a general rule, we would suggest using nonlinear registration whenever aligning volumes between different subjects, since the anatomical structure among even control subjects can vary to a wide degree.

In looking at the BMN comparisons, oddly enough, nonlinear warping to standard space *was* used for one software (SPM) but not for the other two (AFNI and FSL). The reason for this discrepancy is unclear, and in terms of comparisons of final results, this processing difference likely would have a very large effect in the relative ability for voxels from the same physiological region in subject space to be aligned for the final analyses. Also unclear is BMN’s choice of a highly non-optimal reference template, since their selected MNI_avg152T1 dataset is both smooth (low structural contrast) and low resolution (2x2x2 mm^3^ voxels); see Fig. 3, first column. This does not provide a strong target for bringing subject anatomical features into close alignment from ~1x1x1 mm^3^ voxels; Fig. 3, second and third columns show that the average group alignments are generally accurate for the largest structures, but highly variable for finer gyri/sulci.

**Figure 3.**
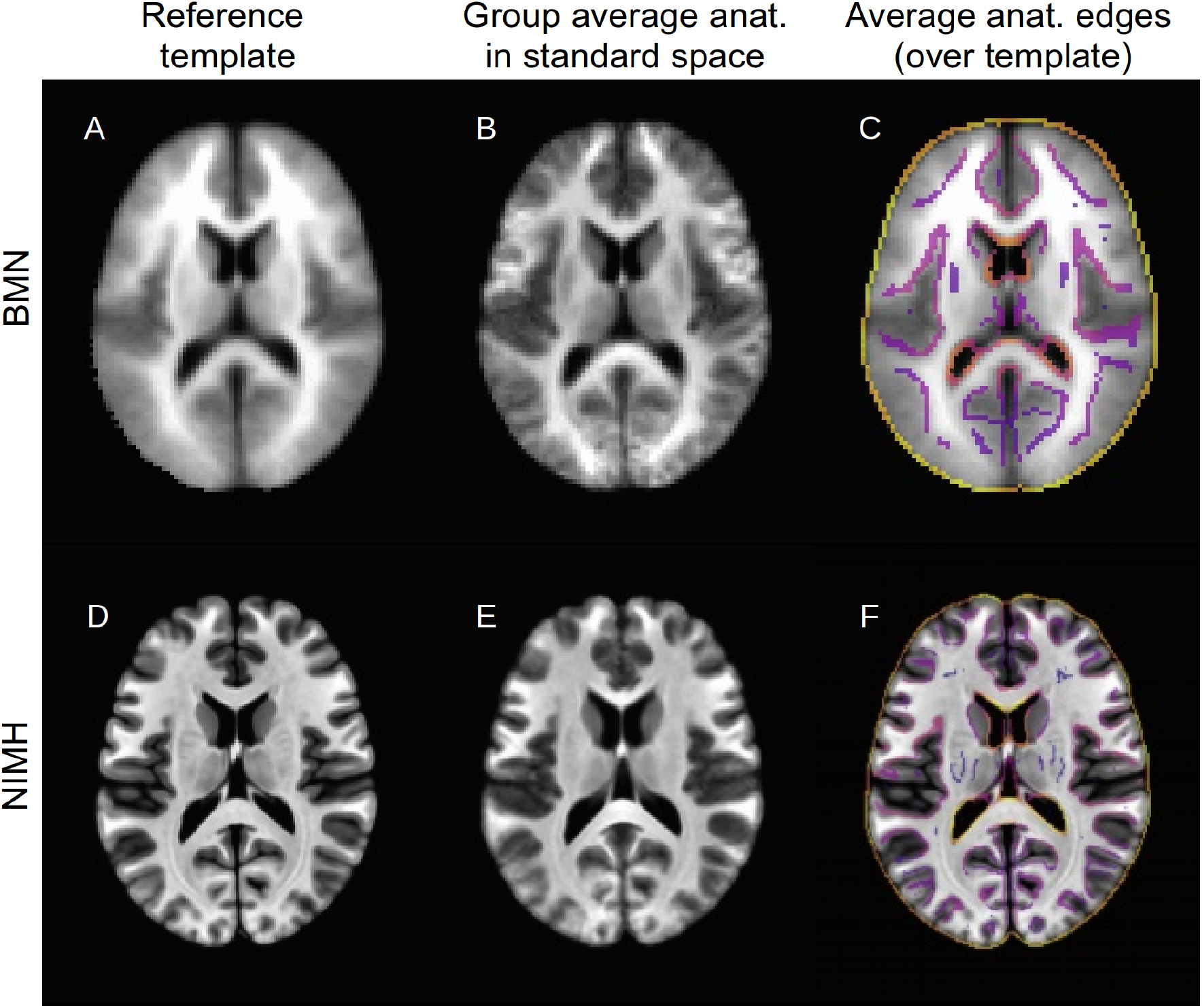
A comparison of the alignment to standard space for BMN- (top row) and NIMH-AFNI (bottom row). The first column shows the target volumes for each processing stream: the blurry and low-resolution MNI_avg152T1 for BMN (A), and the MNI152_2009_template used here (D), which provides more detailed structure for matching features. The second column shows the results of averaging the aligned anatomicals for each processing stream: *anat_final* for BMN (B) and *anatQQ* here (E). Blurry areas show regions of higher variability across the group; while some posterior regions from the NIMH-AFNI stream show noticeable blur, the BMN-AFNI result shows much less spatial contrast there and globally, as expected from the alignment method and template target used. The third column shows the reference anatomical for each processing as a grayscale underlay, and the edge-ified view of the group average anatomicals (via AFNI’s 3dedge3) as plasma-colored overlay. In both cases, the average structures show reasonable alignment to the targets, but the NIMH-AFNI stream (F) has both more features in the anatomical to align to, and more clear features in the results. In all cases: axial slice, *z*=15S, image left=subject left.

Here, we use nonlinear warping of each single subject’s anatomical through the @SSwarper command, which accomplishes the dual task of 1) aligning each subject’s anatomical to a standard space target and 2) skull-stripping that anatomical volume.^10^ The warps created by this script are passed as options of the “tlrc” block in afni_proc.py, as well as the information that the anatomical has been skull-stripped (“-copy_anat DSET” and “-anat_has_skull no”). For the target anatomical, here we choose the MNI152_2009_template volume (see Fig. 3, first column, second row), which has clearly discernible gyral and sulcal features and 1x1x1 mm^3^ voxels, properties that provide stronger features for the warping tool to leverage in reducing the variability of intersubject alignment (even if the final resolution of the EPI data will be lower than the template’s). The average of group anatomicals shows much finer details in Fig. 3 (second and third columns), and the comparison of individual subject alignments in Fig. 4 highlights this further.

**Figure 4.**
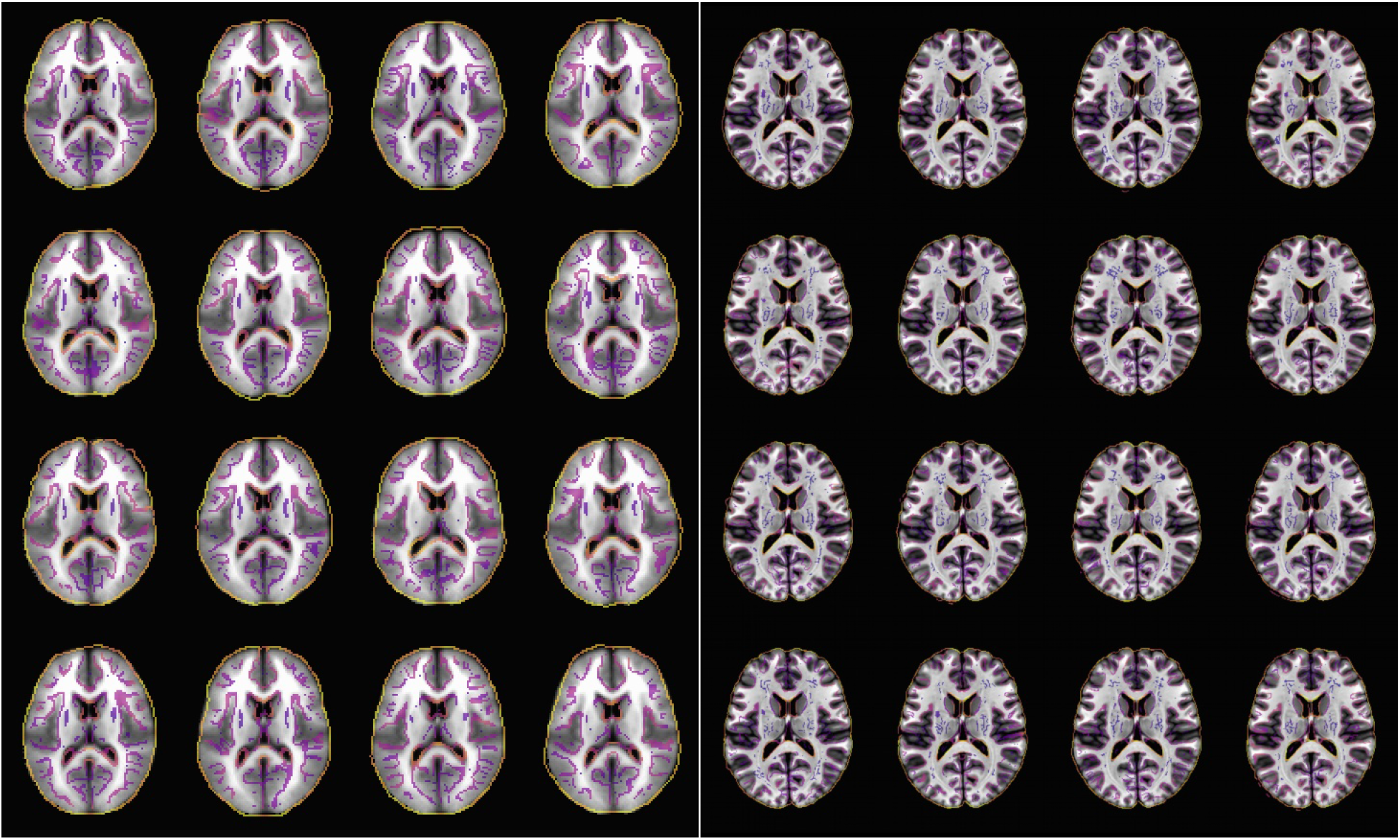
Results showing alignment of each subject’s anatomical to standard space for the linear affine BMN approach (left) and the nonlinear NIMH-AFNI approach (right). The grayscale underlay is the standard space template for each approach, and the plasma overlay is an edgified view of the aligned T1w anatomical for each subject. While alignments show general agreement in all cases, the nonlinear NIMH-AFNI approach tends to show much greater matching of anatomical detail across the brain. In each axial, *z*=15S, image left=subject left.

#### [B-2] Alignment of EPI to anatomical: cost function selection

Aligning functional EPI data to the subject’s own T1w anatomical volume can be difficult because the former typically have low structural contrast and spatial resolution, as well as different relative tissue contrast. To address these issues, the “local Pearson correlation” (lpc) cost function was developed (Saad et al., 2009).

However, even with that cost function, alignment can still fail, due to such properties of the datasets as differing obliquity, relative rotation, and FOV origins recorded in such a way that the volumes may not overlap strongly. Therefore, additional features can be flagged with special options to prealign centers of mass (such as “-align_opts_aea -giant_move”), and BMN did make use of these. However, when *still* facing alignment issues for some subjects (as they note, the EPI datasets here have little structural detail), BMN then utilized the mutual information (“mi”) cost function. Instead, we would recommend trying the “lpc+ZZ” cost function; for most alignments, results should be essentially the same as with “lpc,” but there is an additional stabilizing optimization step that helps in problematic cases. The “lpc+ZZ” cost function is currently the recommended go-to in AFNI when aligning EPI and T1w volumes (or any volumes with inverted contrasts).

An example of alignments using each of the cost functions is shown in Fig. 5, showing edges of the EPI dataset overlayed on the subject’s anatomical (after mapping to standard space, for ease of viewing similar slices). While both cost functions produce reasonable overall alignments, the lpc+ZZ results show more consistent alignment throughout the internal structures and at the posterior edge. For both cost functions, the largest differences between structure and EPI edge appear to be in the anterior regions, and these appear to be due to the presence of EPI distortion, which causes both geometric and signal-value distortions and occurs along the latter part of the phase encoded axis (typically the *y*-axis in the direction of posterior to anterior, matching the locations of observed distortions here). Fig. 6 highlights some of the presence of EPI distortion by comparing the extent of high BOLD signal in two sagittal slices (shown in alignment to standard space) for a single subject.

**Figure 5.**
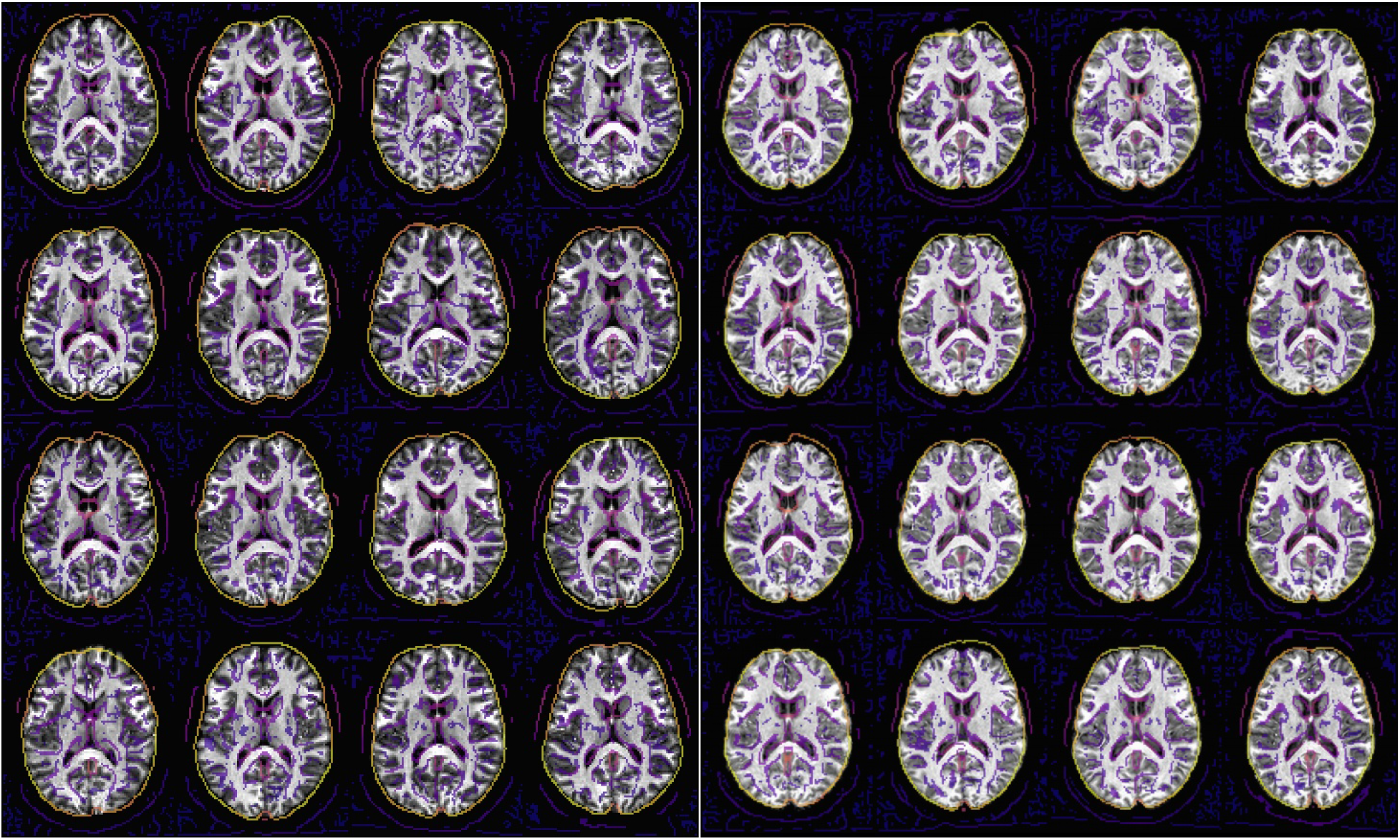
Results showing registration of each subject’s EPI functional data to their own anatomical for the BMN approach (left) and the NIMH-AFNI approach (right). The grayscale underlay is the anatomical of each subject in standard space, and the plasma overlay is an edgified view of the [0]th volume of the run-01 EPI dataset for each subject. While the edges show reasonable fits overall in both cases, those on the right tend to have consistently better alignment with internal and posterior regions in particular; the differences in the anterior regions appear to be due to the presence of EPI distortion. In each axial, *z*=15S, image left=subject left.

**Figure 6.**
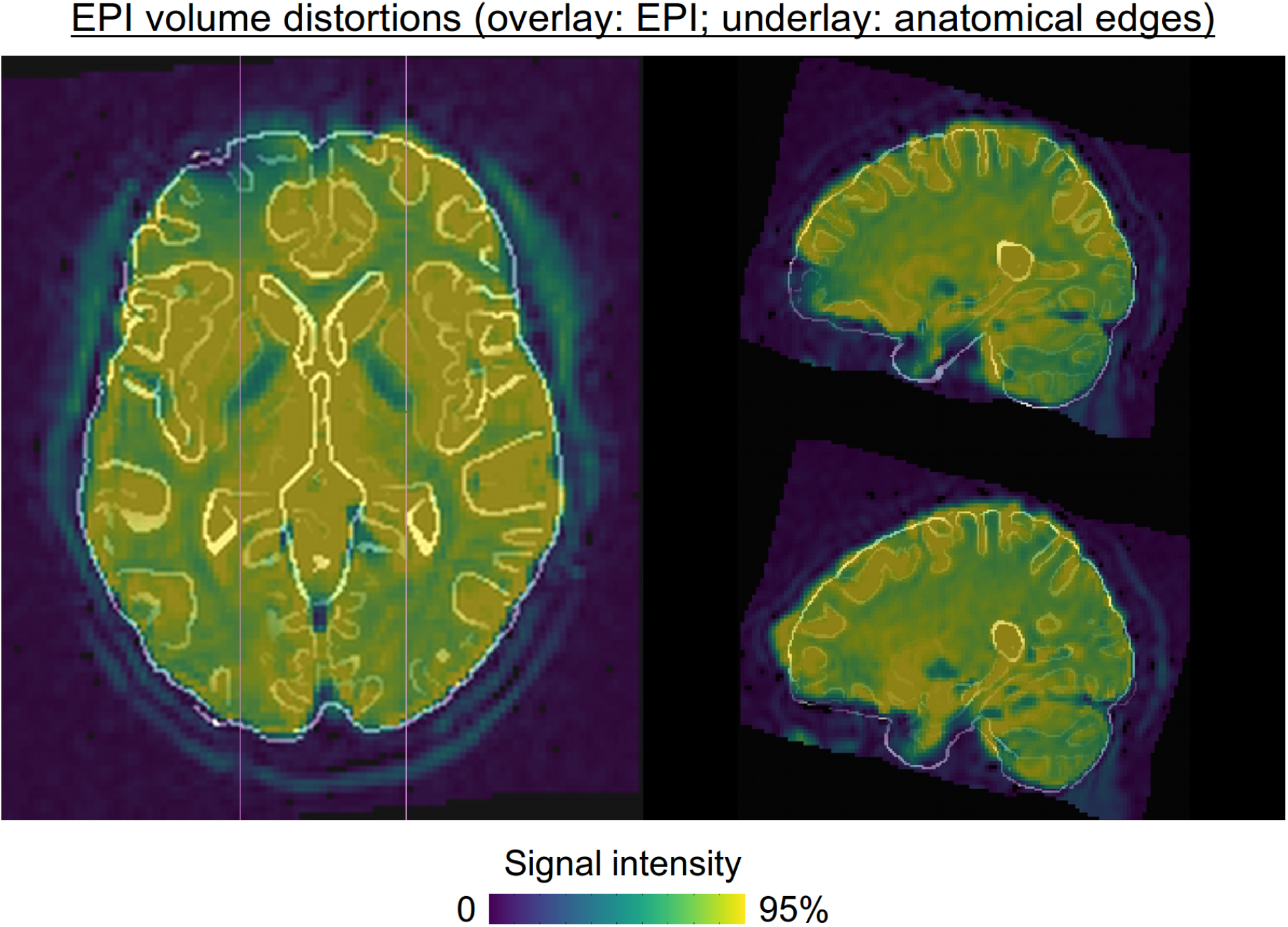
Example of geometric and signal distortions in the functional BOLD data (overlay) due to EPI distortion and signal dropout (sub-02 from ds000001). The subject’s EPI data is overlaid on an edge-ified version of their anatomical (after alignment to standard space; the color scale reflects relative magnitude). The left-right homologous sagittal slices at *x*=25L and 25R show the relative geometric and signal distortions present. The axial slice at *z*=0S also shows the locations of the sagittal slices (magenta). In axial, image left=subject left; in sagittal, image left=subject anterior.

#### [B-3] EPI volume registration for motion correction (and alignment)

When performing volume registration of the EPI time series across time, one volume must be selected as a reference or target of comparison for the others; the relative rotations and translations estimated to register the other time points then provide time series regressors of motion. Additionally, the selected EPI volume is then also used for alignment to the subject’s anatomical volume. Therefore, it is important to avoid selecting a reference EPI volume that is itself corrupted by motion, dropout or other artifacts. Choosing some pre-specified *i*-th volume from the time series does not provide any security or assurance against it containing artifact; one would have to check the acquired data itself in some manner.

In afni_proc.py the recommended way for selecting an EPI reference is to use the following heuristic: select the volume with the fewest number of outliers in the initial EPI time series (denoted by the following option and keyword “-volreg_align_to MIN_OUTLIER”). In general, this has been a quite robust method to avoid having a very poor quality or motion-corrupted volume play the dual reference role.

### C) Further comments on the processing

#### [C-1] The question of upsampling voxel size

A common spatial resolution of acquired functional EPI datasets is a voxel edge length of 3-3.5 mm, with the result that common voxel volumes are ≈27-43 mm^3^. For example, the presently analyzed dataset had 3.125x3.125x4 mm^3^ voxels, with a voxel volume of ≈39 mm^3^. The spatial resolution chosen by BMN to upsample the functional data was 2x2x2 mm^3^, and thus a volume of 8 mm^3^, which represents upsampling by nearly a factor of 5. Presumably, this was chosen because it is the default in the SPM and FSL processing pipelines, though even this “default” is not always followed, such as in some validation tests (see Flandin & Friston, 2017). In the afni_proc.py processing implemented here, we did also use the BMN-chosen final resolution of 2 mm isotropic voxels but solely for the sake of comparison with their results. We note that cluster results and FPR values do not appear to depend strongly on resampling in AFNI (Cox and Taylor, 2017); however, in some processing cases a stronger dependence of outcome on spatial resampling does appear possible (Mueller et al., 2017).

In general, we do *not* recommend upsampling the data by large amounts during processing. One mooted benefit of upsampling might be a more appealing visualization of the data and results in the eyes of some beholders. However, upsampling does *not* create new information at a finer resolution, so no real feature definition is gained, and the upsampling scheme itself may affect the results. It also has the practical downsides of greatly increasing processing time and disk space requirements to hold data, as well as reducing the portability of data (e.g., from processing clusters to local computers, or uploading for data sharing projects). Moreover, radical upsampling further aggravates the multiple testing issues brought about by the large numbers of voxels in the brain. Lastly, excessive upsampling can cause significantly underestimated smoothing in some software implementations, resulting in a higher FPR than the nominal target (Mueller et al., 2017).

We prefer preserving the acquired voxel size and presenting results that closely reflect the spatial resolution of the original data; see, for example, the raw functional and anatomical volumes for one ds000001 subject in Fig. 7, which show the relative coarseness of the acquired data. By default, in afni_proc.py the final voxel grid size will be slightly rounded to a finer matrix than the acquired data; specifically, the value of the minimum edge length, truncated to 3 significant bits. (Note that we recommend researchers to acquire approximately isotropic voxels, as well.)

**Figure 7.**
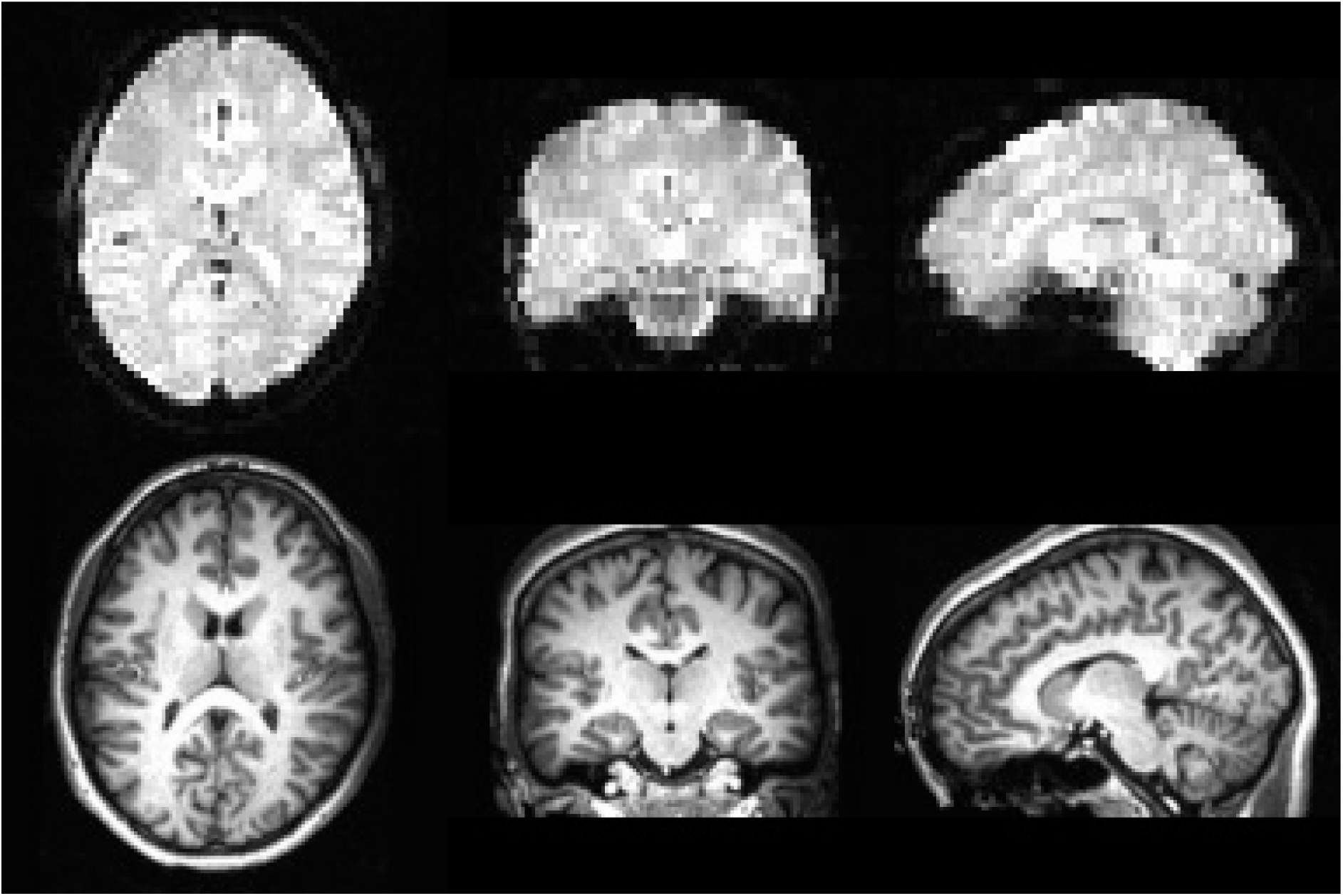
Axial, coronal and sagittal slices of the acquired MRI data, at original resolution and contrast for ds000001 sub-01: functional EPI at 3.125x3.125x4 mm^3^ (top) and T1w anatomical at 1.0x1.3x1.3 mm^3^ (bottom). While upsampling the functional data to 2 mm isotropic voxels will make final result maps look smoother, the process does not add information or detail, which was acquired (as shown) at much lower resolution. In axial and coronal: image left=subject left; in sagittal, image left=subject anterior.

#### [C-2] Specifying statistical tests and selecting voxelwise p-values

BMN utilized voxelwise one-sided *p*-values of 0.01 for thresholding the ds000001 dataset, in order to match the one-sided testing chosen by the original researchers (Schonberg et al., 2012). While voxelwise *p*-value thresholding always carries some arbitrariness with it, this particular value is typically no longer recommended for cluster thresholding based on a preselected voxelwise *p*-value, as FPR control may be more difficult in general (see Eklund et al., 2016; Cox et al. 2017a; Cox et al. 2017b; and this was also noted by BMN themselves). For parametric clustering within AFNI, typically *p*-values of 0.001-0.005 are more commonly used. In the present replication we did implement the unadjusted and large one-sided *p*-value threshold of 0.01 for ds000001, solely to match value used by BMN, which itself was chosen to match the choice by the original researchers (Schonberg et al., 2012).

We further emphasize that just reporting a *p*-value (e.g., as a voxelwise threshold for clustering, or for any usage) does not provide enough information to the reader (see A-4, above). The investigator *must* also report whether one- or two-sided testing is being performed, and in the particular case of choosing one-sided testing, specific justification *should be* provided, otherwise a two-side testing should be adopted (or equivalently, one should choose a voxel-wise one-sided *p*-threshold that is half of the intended value). BMN set up their voxelwise analysis as a pair of one-sided tests, looking for positive and negative results separately. As noted above, simultaneously having two separate tests necessitates a multiple comparisons adjustment factor of 2; that is, if a two-sided test were performed with threshold voxelwise *p*=0.01, then each of the paired one-sample tests, as implemented in BMN, should be thresholded with voxelwise *p*=0.005 for equivalency.

Finally, in order to further reduce the dependence on the arbitrary choice of a single voxelwise *p*-value threshold, there is a new tool within AFNI with a more “equitable” dependence on such parameters for clustering (Cox et al. 2017a; Cox, 2018). It is important to note that use of this “ETAC” clustering paradigm would dictate some specific processing choices when setting up afni_proc.py, like not blurring data during pre-processing. More description and implementation details are described within the aforementioned sources or via the AFNI Message Board.

#### [C-3] Presenting results: including the effect estimates

We would also present beta coefficients as voxelwise effect estimates. These should be meaningful, and then can help QC the results for “sanity checks” (Chen et al. 2017). In addition, reporting effect estimates can promote reproducibility and assist more accurate meta analysis than the current coordinates-based approach. Note that different software packages perform scaling in very different ways during preprocessing, and this may lead to different interpretations of effect estimate and group analysis accuracy. In AFNI, the time series are scaled by the voxelwise mean, so that the effect estimates can be more meaningfully interpreted as an approximation for BOLD percent signal change at each voxel (Chen et al. 2017), rather than the acquired (unitless) data magnitude. In AFNI, we approach this in a localized (i.e., voxelwise) fashion and therefore the values are directly comparable across brain regions and subjects at the group analysis stage; other software calibrate in a “global” manner using brainwide averages, which produces different results and interpretations. The effect estimates form the basis of the group level modeling, and we feel strongly that the values should be presented in images, since they convey valuable information as the primary quantity of interest; the complementary statistic information^11^ is typically represented through thresholding. In any area of science, it is important to view and display the main quantities being estimated.

Even though BMN did not explicitly mention this step in processing, they did include the “scale” block in their afni_proc.py command; this is something we do recommend, and it allows for interpreting the time series and effect estimates as percent signal change. Fig. 8 shows the effect estimate maps for ds000109, corresponding to the bottom row of Fig. 1. The maximum magnitude of effect in this case was approximately 0.5% BOLD signal change (and the same voxelwise thresholding is carried out via the statistic values).

**Figure 8.**
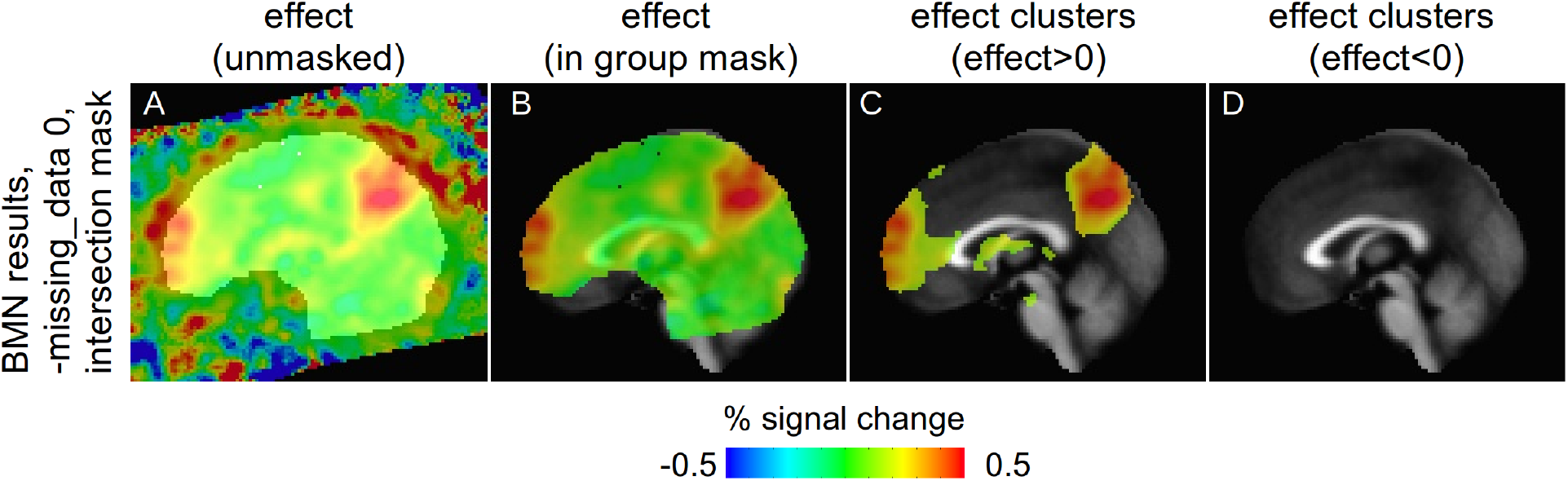
Effect estimate maps for ds000109, corresponding to the bottom row of Fig. 1. We generally recommend using the “scale” block in afni_proc.py to implement per voxel scaling the EPI data, after which the effect estimates are interpretable as approximations of BOLD percent signal change relative to the baseline (as is the case here). These effect estimates form the basis of statistical testing, and we feel strongly that they should be shown when presenting results (voxelwise thresholding is still done according to the statistic values, as in panels C and D; as noted in Fig. 1, this was done at *p*=0.005 and FWE=5%, leading to a cluster size threshold of 442 voxels here).

#### [C-4] The question of removing initial EPI volumes

BMN removed the first two TRs of the EPI functional time series in their processing. Initial TRs are often removed in early stages of processing in this way when they contain signals from pre-steady state magnetization. However, in the present data that does not appear to be the case (perhaps, as some scanners are configured, pre-steady state volumes had *already* been removed), so that the amount of data here need not be reduced. In our analyses here, we also removed the TRs in order to match BMN’s analysis; however, we likely would not have elected to remove the volumes, as there were stimulus events over that interval. We note that, in the scenario of pre-steady state volumes existing in and not being removed from processing, afni_proc.py would likely detect this and warn the user (via the @ss_review_* scripts).

We note that it is often beneficial *not* to remove pre-steady state TRs from time series prior to using afni_proc.py. Those volumes can actually be useful for registration, because they often have higher tissue contrast than the post-steady state TRs, and therefore can be very useful as a reference volume for alignment to the subject’s anatomical (Gonzalez-Castillo et al., 2013); such volumes would be removed at the onset of afni_proc.py, and aside from the possible use in registration, would have no impact on the analysis.

#### [C-5] Yes, afni_proc.py can do motion-based censoring

BMN make the following comment in Section “2.2.4 ds000109 Analyses” (emphasis added):

> … *custom software was applied to exclude functional volumes where head motion had exceeded a certain limit, however this process was omitted from our pipelines **since this feature was not available in any of the software packages***.

This is surprising and confusing to read, because: 1) AFNI does have the capability to censor volumes based on head motion estimates (since 1998, in fact), and 2) BMN themselves did implement the option to censor data in their afni_proc.py command (as “-regress_censor_motion 0.3”). We note that the use of this option for motion-based censoring is, indeed, one that we recommend using in standard processing.^12^

Additionally, we often suggest also adding another afni_proc.py option to remove volumes that are likely “bad”, based on where too many voxels are flagged as being outliers (by 3dToutcount). This can be implemented in afni_proc.py by adding the option “-regress_censor_outliers LIMIT” where LIMIT is a fraction like 0.1 or 0.05.

#### [C-6] Motion regression in the presence of multiple runs

To better account for motion, we recommend applying -regress_motion_per_run in afni_proc.py. As the name suggests, it causes motion parameters to be modeled separately per run.

#### [C-7] Automatic single subject and group level QC with afni_proc.py

Part of basic processing includes checking the results of each stage and evaluating whether problems have arisen. This includes looking at the quality of the various alignment steps, assessing motion, checking any collinearity or modeling problems, and counting remaining degrees of freedom (DOF) after regression.

At the single subject level, afni_proc.py automatically calculates and stores several representative quantities about the datasets (for example, in both the BMN and present processing). These are displayable with @ss_review_basic (Fig. 9) and @ss_review_driver (Fig. 10), the latter of which also opens up AFNI interfaces to display volumetric and other data (such as motion parameters, outlier counts, etc.). Thus, researchers may directly evaluate the amount of motion, quality of alignment, and other important features in a given processing stream.

**Figure 9.**
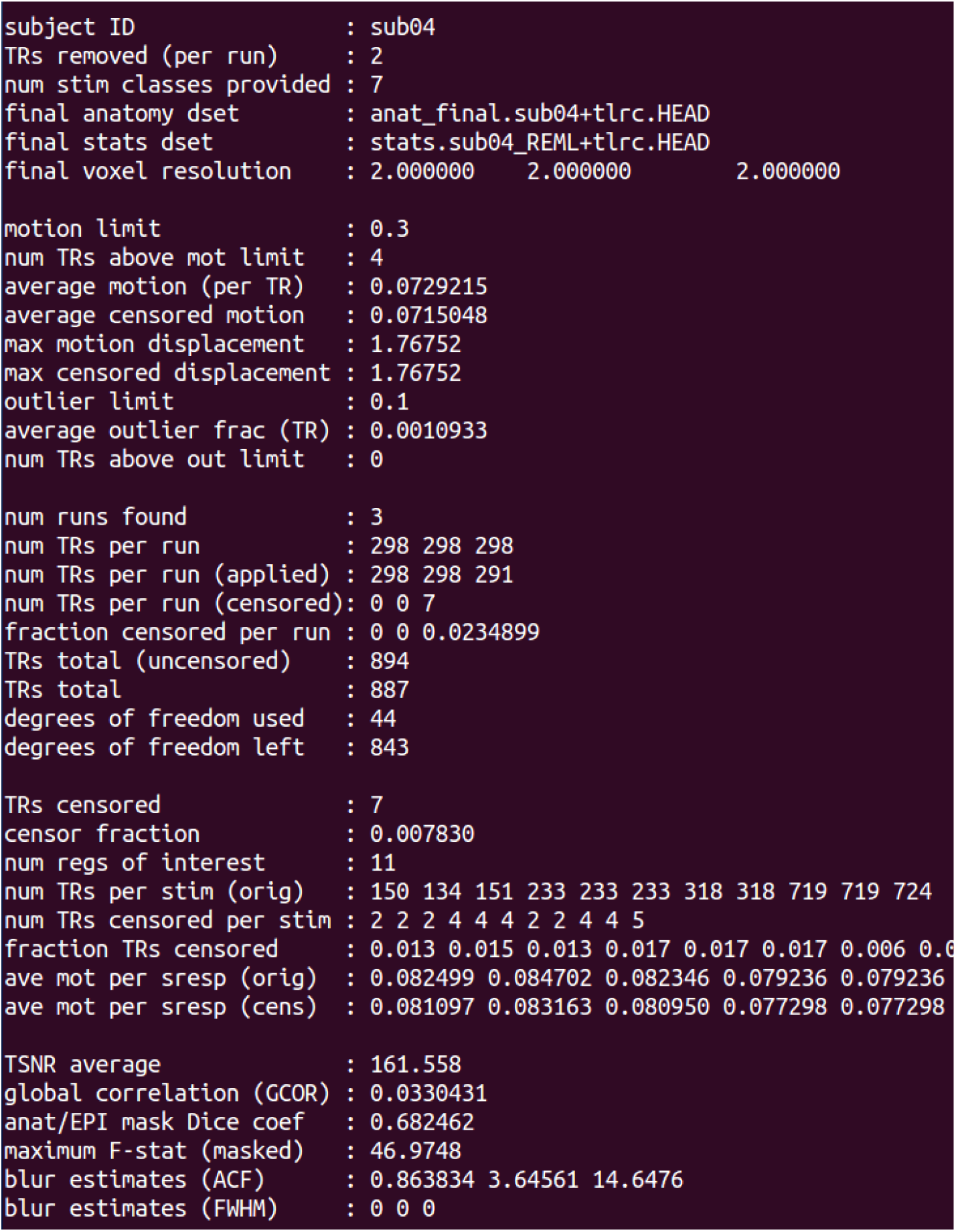
For single subject QC, descriptive quantities automatically calculated and stored for each subject when using afni_proc.py (here, displayed using @ss_review_basic). At the group level the AFNI program gen_ss_review_table.py can combine these outputs into spreadsheet table for summarizing group properties and efficiently detecting outliers (see Fig. 11).

**Figure 10.**
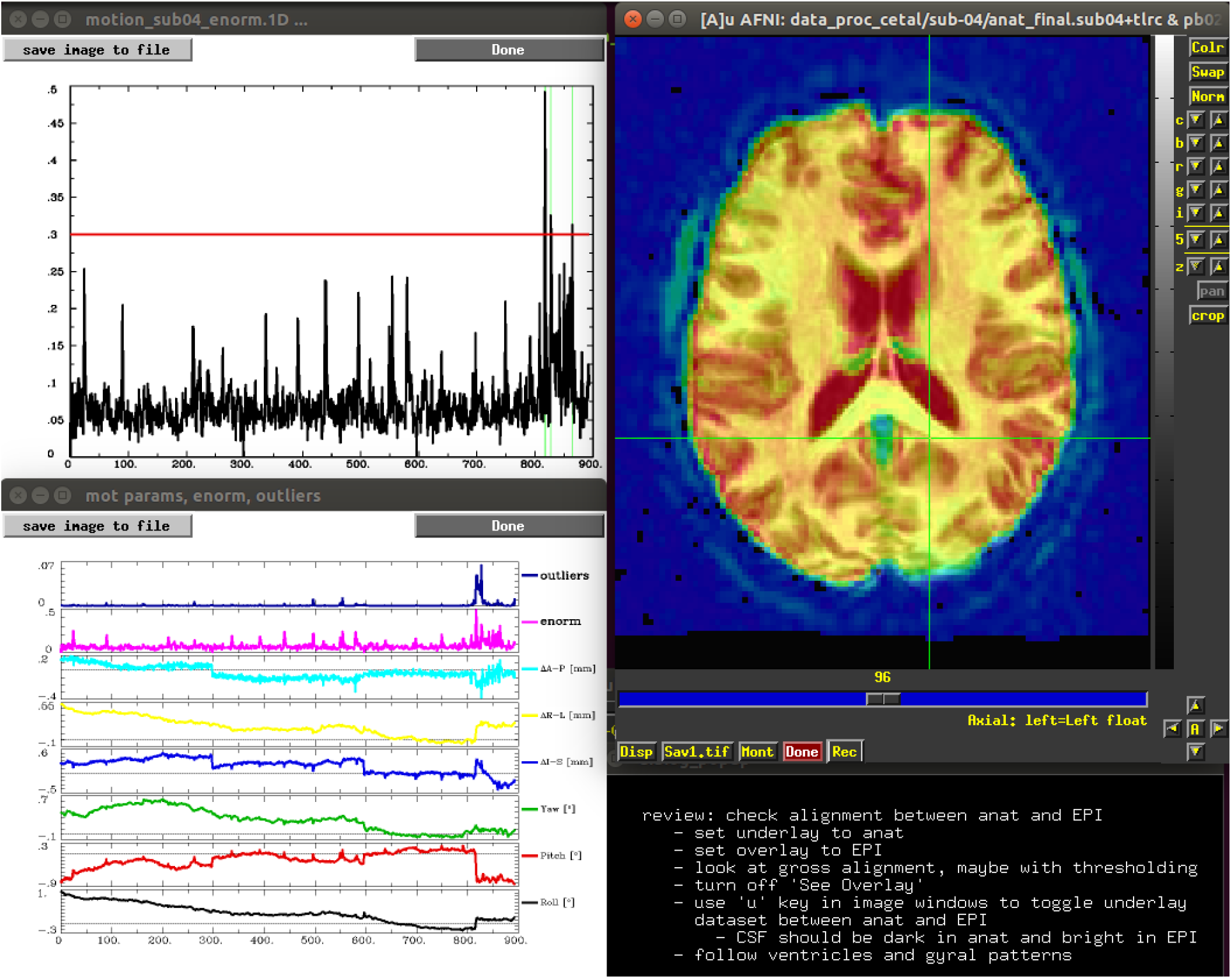
A screenshot of part of the guided QC output using @ss_review_driver, a script which is created by afni_proc.py for each subject automatically. Exhibited data includes: motion parameter estimates, Euclidean norm of motion and outlier counts over time; the Euclidean norm of motion with the user-defined threshold in red as a reference; and volumetric data sets overlaid to check EPI-to-anatomical registration. A pop-up dialogue screen provides guided QC suggestions.

At the group level, the AFNI program gen_ss_review_table.py can be run to combine all the single subject review outputs into a single spreadsheet table (Fig. 11). For example, with the command:

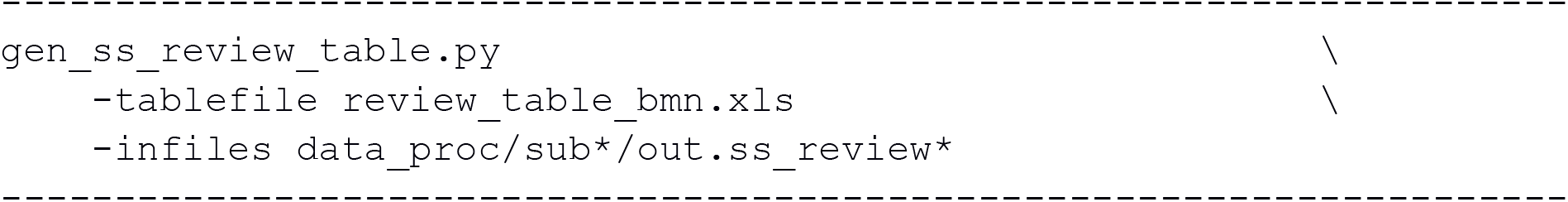

**Figure 11.**
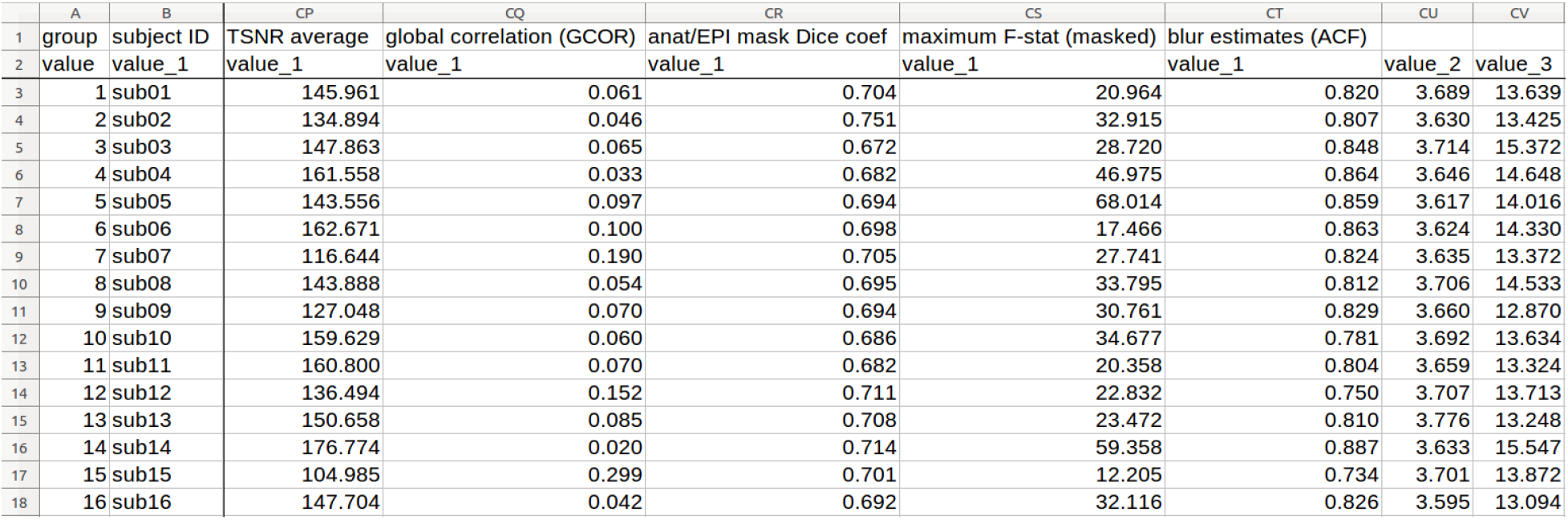
A partial view of the group level table created by gen_ss_review_table.py, which combines useful metrics from the individual subject processing for easy comparison and analysis. This aids in summarizing group properties and detecting outliers.

From the combined results, one can directly evaluate the distributions and ranges of relevant properties, as well as detect outliers (e.g., in terms of “average censored motion” or “censor fraction”). For example, from the NIMH-AFNI processing, it is possible to tell quickly that this group’s TSNR had a mean±standard deviation of 145.1±18.9 and a [min, max] = [105.0, 176.8]; one could use this to evaluate the scanner protocol and acquisition.

#### [C-8] Making a 3dMEMA command efficiently

Scripting can be difficult and time consuming: getting paths, filenames, volume selection and other features correct, even when “just” adjusting existing scripts to other groups/studies. The 3dMEMA command call can potentially be quite lengthy, because one inputs multiple volumes per subject, and it typically involves also including labels and volume selectors. However, with reasonable organization of file structure, this procedure can be greatly simplified through the use of AFNI’s gen_group_command.py program.

Using the present dataset as an example, instead of writing a 3dMEMA command directly with 16x2 arguments of long file names (16 subjects, each having both a beta coefficient and a t-statistic volume to include), as well as other options, gen_group_command.py can accomplish this with 3 options (plus the other control options):

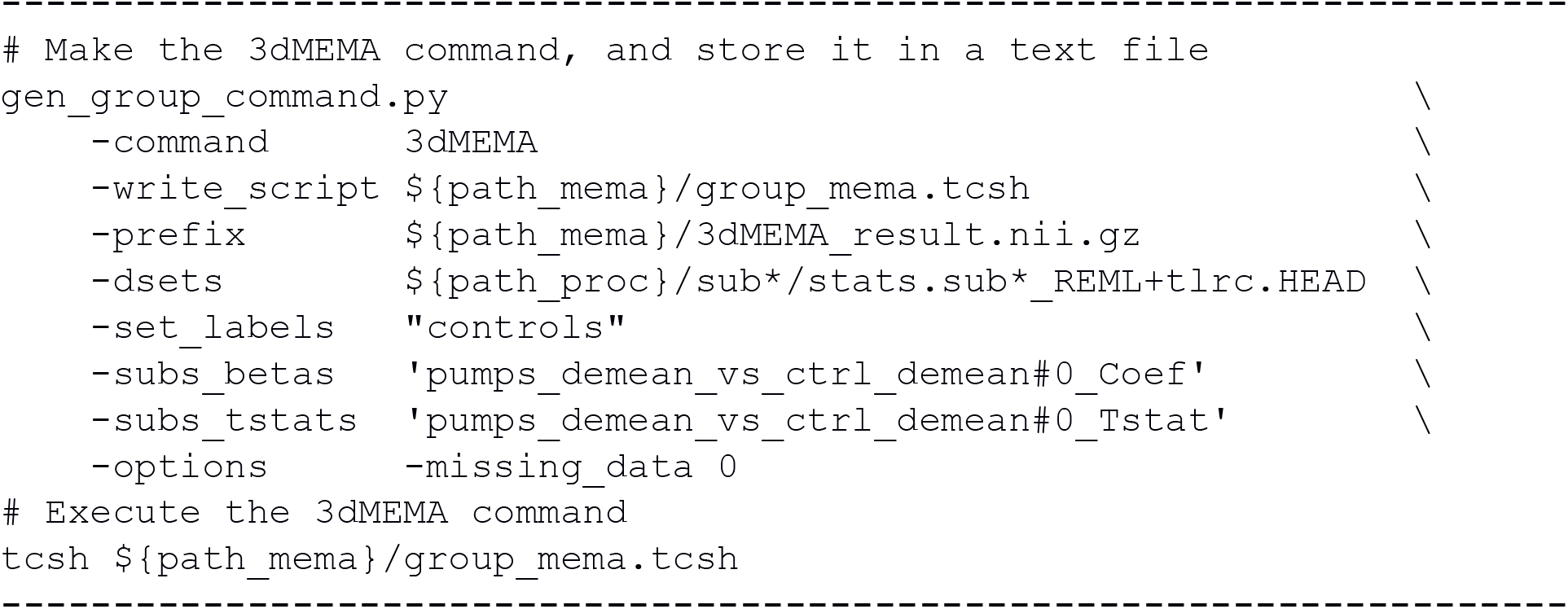

Note that this generator program becomes even more efficient to use as the number of subjects grows; the constructed 3dMEMA command becomes much longer, but the syntax here would remain unchanged.

#### [C-9] Output format

As a final processing comment, we note that AFNI programs can output NIFTI standard volumes directly. Typically, this is done by specifying the output file name with a “.nii” or “.nii.gz” suffix, such as “-prefix OUTPUT.nii.gz.” A separate conversion step is not necessary in most cases. While the afni_proc.py pipeline does output only BRIK/HEAD-formatted files, the remaining programs’ outputs (such as clusters, supplementary masks, etc.) can be output in NIFTI format directly.

### D) Brief comparison of BMN- and NIMH-AFNI processing results for ds000001

Having described several differences in individual processing steps between the BMN-AFNI and recommended NIMH-AFNI processing streams, we now briefly show the cluster results of each for ds000001, following the presentation of BMN-Figure 1 (first column). Fig. 12 presents the groupwise *t*-statistic maps for the whole FOV and brain mask, as well as the separate “positive” and “negative” cluster results from the separate one-tailed *t*-tests are shown, for the separate cases of BMN processing (with and without the inclusion of the zero-masking feature mentioned above in A-1) and the NIMH-AFNI steps. The whole brain mask volumes for BMN- and NIMH-AFNI were 245,683 and 220,281 voxels, respectively.

**Figure 12.**
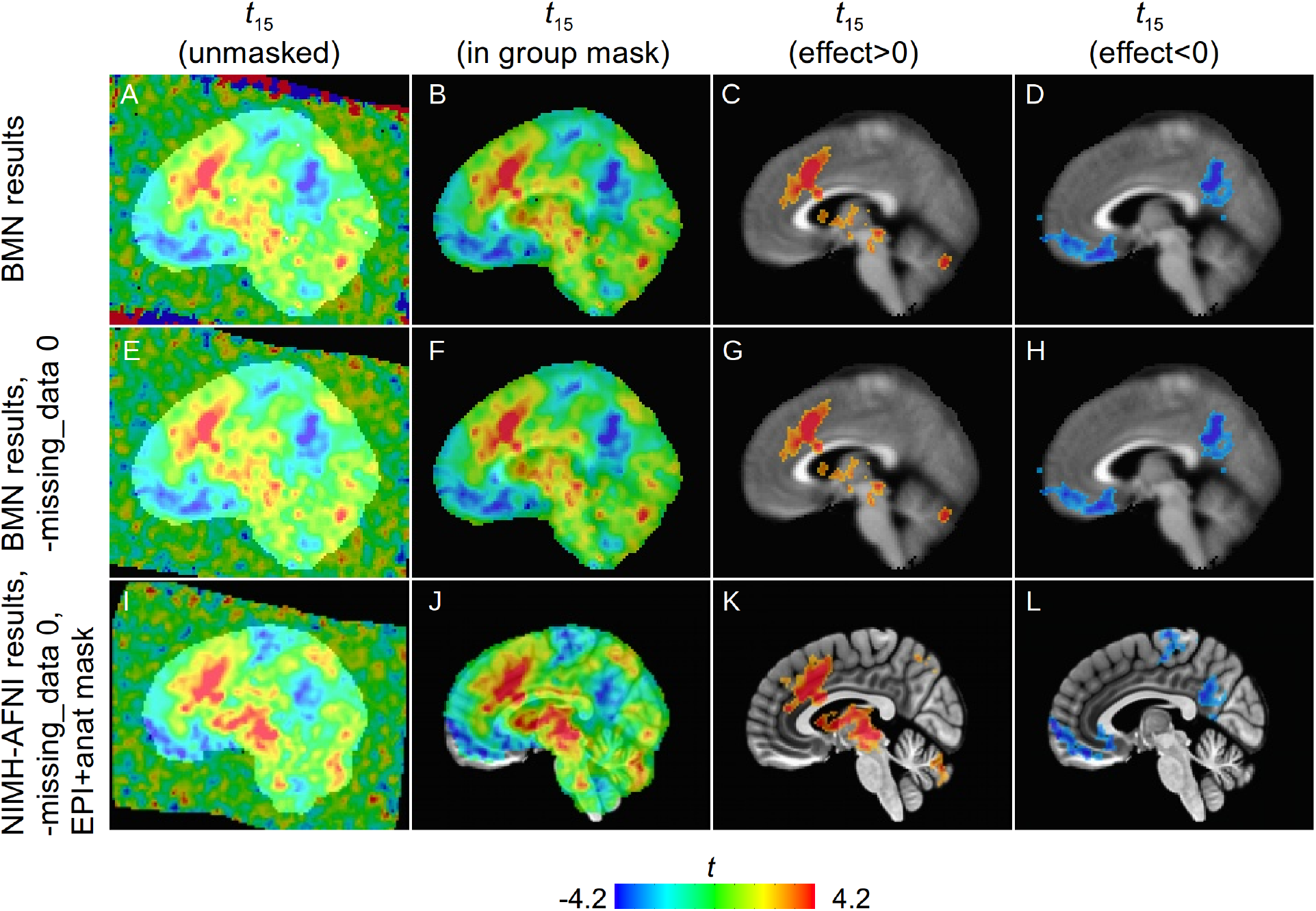
A comparison of processing, masking, group level approaches with ds000001 (cf BMN Fig. 1, left column). In the top row, A shows BMN’s unmasked *t*-statistic maps (overlaid on their group mask), with very large magnitudes (~100 in most cases), due to their inclusion of edge artifacts by not masking out zero-values (see Fig. 1 caption and text description A-1); in this case these edge-artifacts do not occur within the BMN group mask, though that mask is still likely too large. In the middle row (E-H) 3dMEMA was run using the correct “-missing_data 0” option, and the artifacts are not present. In the bottom row (I-L) we display the results of the NIMH-AFNI processing stream, including its group-level mask (underlaid in panel I), which is created from each subject’s anatomical and EPI data rather than from the template extents (discussed more in point A-2). Effect estimates of this data are displayed in Fig. 13. In all cases: sagittal slices; *x*=4R; image left=subject anterior.

**Figure 13.**
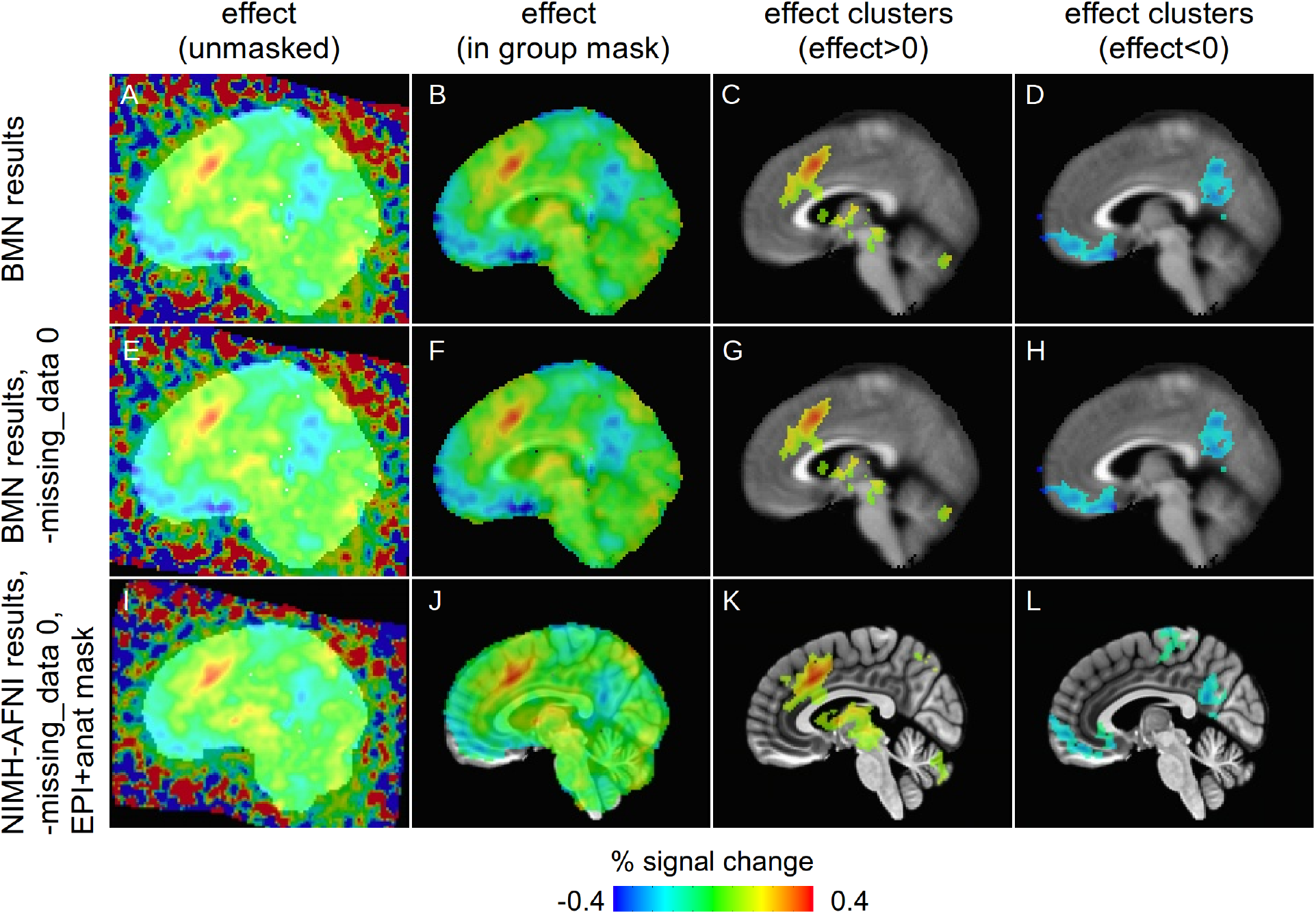
A comparison of group level results for ds000109 in terms of effect estimates (with AFNI’s scaling, these represent BOLD percent signal change). Statistic value information is used for voxelwise thresholding in the last two columns. See Fig. 12 for descriptions of each panel. In all cases: sagittal slices; *x*=4R; image left=subject anterior.

There are many similar clusters output from the BMN and NIMH-AFNI processing, with some differences noticeable in cluster extent (minimum cluster threshold volumes of 309 and 242 voxels, respectively; (voxelwise threshold *p*=0.01, FWE=5%, one-sided test, NN=1) and location. Without a gold standard for comparison, it is difficult (if not impossible) to compare these clustering results in a more meaningful manner.

We also note that, based on the @ss_review_basic and gen_ss_review_table.py results (see C-7, above), we would also likely recommend removing sub-15 from the group analysis as an outlier. The group’s mean±stdev for “average outlier frac” is 0.0022±0.0049, with the value for sub-15 being 0.0204 (and the next largest is 0.0024); additionally, that subject had >30% of volumes censored in each run. Leaving in such subjects may be considered an additional source of noise or artifact.

## DISCUSSION AND CONCLUSION

We have outlined here several considerations for researchers when analyzing FMRI data. For this particular set of data, we have described specific recommendations for initial processing using AFNI and its primary tool for designing a customizable (yet also understandable) processing stream, afni_proc.py. Supplementary AFNI programs and approaches for performing nonlinear alignment, data visualization, single subject processing QC, and group property QC have also been presented.

Many of the considerations discussed here involve the question of default processing steps. Technically, a “default” setting is one used by a program when no option is specified; oftentimes, these are assumed to be the pre-selected “go-tos” or “best initial settings” for that particular program. However, in AFNI this is not the case, due to the combination of providing continual development while also maintaining backward compatibility. Changing even a single program’s default (in the technical sense) can potentially lead to changes in results without a researcher’s knowledge. Changes to defaults must be done with care and should be well-advertised (which we do try to do when these changes to become necessary). Therefore, in the context of AFNI, we typically update recommendations and examples rather than default program settings.

One common feature of processing described here, and a central focus of AFNI, is to visualize one’s data carefully and often during processing. To repeat a statement made above: QC *is* part of processing itself. This is one reason why we do not recommend deleting intermediate data; while this practice may take up more storage space, it also permits one to visually check steps for each subject such as alignment, masking and outlier evaluation, each of which has a tremendous impact on the final outcome. As noted by BMN with some of the OpenfMRI sets analyzed here, data quality and contrast can lead to the need for re-processing and to the selection of different options or parameters.

Additionally, even seasoned analysts may make mistakes in data processing (e.g., typographic errors, incorrect filename specification, or misselecting an option), and looking at data can provide the best method for detecting them. As an example from the present study, the cluster of *t*-statistics in the ds000109 results in the cerebellum with values of ±100 reported by BMN were purely artifactual from the presence of the FOV edge; using the appropriate option in 3dMEMA to mask out zero data would have prevented that erroneous result (see A-1), and using a group mask related to the actual data coverage of all subjects would also have helped (see A-2). In the case of being faced with such oddly extreme values, visualizing either the full *t*-statistic image or the overlay of EPI data on anatomical volumes would have provided necessary information to fix this behavior (or at least to generate a question for the AFNI Message Board).

Several features of the afni_proc.py-generated pipelines actively assist in QC and data assessment. The @ss_review* scripts provide counts of outlier volumes, output degrees of freedom, and display images of motion regressors and volumes for alignment. Additionally, numbers such as the TSNR in a dataset (calculated by afni_proc.py and reported with @ss_review_driver) may be a useful check on the acquisition protocol itself: from using gen_ss_review_table.py, it was straightforward to calculate that the TSNR statistics for the ds00001 dataset were 145.1±18.3. As public datasets become more common, knowing such basic properties of the data itself becomes more important.

Reporting processing steps is also an important feature of research. Again, it is worth noting that BMN provided all of their processing scripts. While textual descriptions are useful, especially for providing motivations of choices, only the processing script itself is an accurate record of processing for others to follow: there are simply too many options to list, and longform descriptions may be imprecise. Moreover, simple mistakes may occur only in the descriptions (e.g., typographic errors, forgetting an option, making changes in reprocessing), but having the script itself greatly reduces the effect of such errors. For example, BMN reported in their Methods section that motion-based censoring was not used in their processing, but their afni_proc.py script showed it had, in fact, been used; additionally, their afni_proc.py command showed they also performed time series scaling, even though it was not stated in their Methods. In AFNI, reporting the processing is simplified by being able to include the relatively short (~20-30 line) afni_proc.py command used to generate the processing stream; along with AFNI version number then, there is enough information to reproduce the full script, and the command itself can quickly be viewed to determine primary processing options that were selected.

When reporting results, a large amount of clarity is needed as well. For example, when reporting clustering conditions, it is *not* enough to say, “Data were thresholded at *p*=*X* and FWE rate of *Y*.” As noted in C-2, one must also specify the sidedness of testing, because two-sided testing with voxelwise threshold *X* would be equivalent to a pair of one-sided tests with a threshold of *X/2*. This step may also require restating the specific hypothesis being investigated, to show how it matches statistical testing. It would also be useful to report the group mask size (e.g., voxel count), since this strongly determines the FWE correction, along with the cluster threshold size. We also do not recommend greatly upsampling the FMRI data, as this does not add real information or detail, and does not reflect the reality of the acquired data resolution. To fully describe the results of the modeling, the effect estimates themselves should be presented, as well, since those are the basic quantity of comparison; the complementary statistic information is useful for thresholding or should be shown in separate maps (and should always be accompanied by the degrees of of freedom).

As noted at the beginning, the combined diversities of data type, specifications, available supplementary information, and ultimate research questions make using a pre-packaged pipeline potentially undesirable, though we provide starting points for various analysis configurations (e.g., see the examples in “afni_proc.py -help”). For example, we would generally recommend nonlinear alignment of anatomical datasets among different subjects (including warps to standard space); not upsampling by large amounts; and scaling task time series, unless there are particular constraints for a study. In the present work we have provided several such concrete suggestions for how we would initially approach analyzing datasets such as those provided here. Perhaps more importantly, we have also emphasized the motivations that led to these specific processing configurations -- hopefully these more abstract considerations can be applied generally.

While FMRI processing is complicated and involves many detailed stages of analysis, the inclusion of coordinated quality control as part of the analysis greatly improves the reliability of the final results. We have highlighted several such tools to visualize and quantitatively evaluate processing features within AFNI and particularly with afni_proc.py. We hope that these features are of use to the neuroimaging community and facilitate their scientific studies.

## ACKNOWLEDGMENTS

Thanks go to Tom Nichols for clarifying some points mentioned in the BMN article, as well as some processing features in other software. The research and writing of the article were supported by the NIMH and NINDS Intramural Research Programs (ZICMH002888) of the NIH/DHHS, USA. This work extensively utilized the computational resources of the NIH HPC Biowulf cluster (http://hpc.nih.gov).

## Footnotes

1 As noted in Cox. et al. 2017, scripts can be downloaded from: https://afhi.nimh.nih.gov/pub/dist/tgz/Scripts.Clustering.2017A.tgz

2 Original version, uploaded to bioRxiv on March 20, 2018 (in case revisions occur later). Link: https://www.biorxiv.org/content/early/2018/03/20/285585

3 The name of this modeling differs across software packages. In AFNI this is called “mixed effects multilevel modeling” (MEMA), which is performed using 3dMEMA (Chen et al. 2012).

4 In fact, we would consider QC of each processed dataset to be *part of* the processing steps directly, and not something separate.

5 The NIMH-AFNI and BMN-AFNI processing scripts for ds000001 used here are available at: https://afm.nimh.nih.gov/pub/dist/doc/htmldoc/codex/fmri_brief.html

6 In some cases using a mask restricted to gray matter or at least a higher fraction of gray matter has been suggested for reducing multiple comparisons in a reasonable manner; for example, see Cox et al. (2017a).

7 Mechanically, this could be accomplished with AFNI commands via: *3dmask_tool -prefix mask -input ‘Is ${TOPDIR}/sub*.results/mask_epi_anat. *.HEAD’ -frac 1.0* where *${TOPDIR}* is path to the directory containing all single subject afni_proc.py results. Alternatively, one could make a group mask from the *union* or even fractional coverage of the mask_epi_anat* data sets, as long as the missing data flag is used in 3dMEMA, 3dttest++, etc.

8 Unlike many standard group analysis tests (like a *t-test*), MEMA modeling does not assume that within-subject variance is constant across a group (or much less than the cross-subject variance). In addition outliers can be handled through an optional assumption of Laplacian distribution. It is also worth noting that the cross-subject variance *τ*^2^ is automatically provided in the output from 3dMEMA. For more details, see Chen et al. (2012).

9 On a technical note, BMN correctly applied amplitude modulation via AM2 using the -regress_stim_types option. The two resulting betas (average and modulated response) are stored under the same label, and in order to be specified individually in a symbolic GLT, an index selector must be used, such as ‘[0] ‘.

10 As noted in the OpenfMRI webpage for ds000001, sub-04’s included anatomical was pre-skullstripped; in the BMN approach of skullstripping each anatomical within afni_proc.py, this would have to be flagged separately for this subject (otherwise risking over-clipping the brain). While that is not “wrong”, in the present case where skull-stripping is performed by @SSwarper through alignment with respect to the reference template, the need for separate treatment does not exist -- if no skull sticks out, then the volume would not be pared down further. This is an additional, positive side effect of the present processing.

11 It should also be noted that when reporting statistics (e.g., the *t*-statistic), the number of degrees of freedom (DOF) must be reported as well. In AFNI programs this information is typically stored automatically in the output volume’s header and may be viewed with 3dinfo, for example. For voxels that include some missing data (and would therefore have reduced DOF), the output statistic is transformed to the equivalent *t*-value with the “full” DOF to match voxels with no missing data.

12 From correspondence with the BMN authors, it appears that the use of the word “any” there was a typographic error; they meant to state that: *since such motion-based censoring was not available in* all *packages, it would not be implemented*. However, to avoid confusion over AFNI processing, we feel it is worth pointing out the existence of this option in afni_proc.py.

